# Determining the off-target activity of antibiotics and novel translation initiation sites in mitochondria

**DOI:** 10.1101/2024.08.29.610400

**Authors:** James Marks, Emma Young, Markus Hafner

**Author notes:** Department of Biochemistry, Vanderbilt University School of Medicine, Nashville, TN 37237. Department of Biological Anthropology, University of California, Los Angeles, CA 90095.

## Abstract

The 13 mtDNA-encoded proteins are synthesized using a dedicated translation system that is more similar to bacterial systems than the cytoplasmic system. Consequently, many bacterial protein synthesis inhibitors, used as antibiotics, exhibit mitochondrial toxicity as off-target effects. However, whether these antibiotics act through the same mechanisms in mitochondria as in bacteria remains unclear. To address this, we characterized the impact of a panel of bacterial translation and elongation inhibitors on mitochondrial translation through mitoribosome profiling. We found that the mechanism of action for every antibiotic, except telithromycin, is the same in both bacteria and mitochondria. Additionally, mitoribosome profiling data showed that *MT-ND1* and *MT-ND5* have incorrectly annotated translation initiation sites and suggested the presence of several translation initiation sites on ncRNAs that produce mitoribosome footprints, as indicated by the detection of mitoribosome footprints at these locations. This work demonstrates how antibiotics can inhibit mitochondrial translation by mechanisms identically or very similar to those found in bacteria and the utility of mitoribosome profiling for annotating mitochondrial genes.

## Introduction

Mitochondria contain their own genome, which in mammals canonically encodes thirteen proteins. These proteins are essential building blocks of complexes I, III, and IV of the electron transport chain, as well as ATP synthetase. They are synthesized at the mitochondrial matrix by a dedicated translation system that is conserved from the ancestral bacteria that evolved into the present-day organelles through an endosymbiotic event^1^. As a result, mitochondrial translation is wholly distinct from cytoplasmic translation and utilizes its own mitochondrial ribosomes (mitoribosomes), tRNAs, and translation factors. The RNA components of the machinery are mitochondrially encoded, while its protein components are encoded in the nucleus, translated by the cytoplasmic ribosome, and subsequently imported into the mitochondria^2^.

Chronic impairment of mitochondrial translation, most commonly through mutations in translation factors, results in disrupted mitochondrial function and causes diseases such as MERRF (Myoclonic Epilepsy with Ragged-Red Fibers)^3^, MELAS (Mitochondrial Encephalopathy, Lactic Acidosis, and Stroke-like episodes)^4^, and maternally inherited deafness^5,6^. Furthermore, acute inhibition of mitochondrial translation caused by the off-target action of antibiotics targeting the bacterial ribosome can also cause mitochondrial toxicity^7–10^. This toxicity can manifest as lactic acidosis, ototoxicity^11^, nephrotoxicity^12^, and peripheral neuropathy^13^, depending on the specific antibiotic and the genetic background of the recipient^14^. While the United States has worked to regulate antibiotics with severe side effects, they remain a global issue. For example, chloramphenicol, whose use is discontinued in the United States, causes fatal aplastic anemia at a rate of 1 in 60,000^7^ but remains widely prescribed in low-income countries due to its low production cost^15,16^. Surveys suggest that approximately 14.3 doses of antibiotics are administered per 1,000 people every day worldwide^17^. The toxic effects of antibiotics can be exacerbated when taken without medical supervision. In addition, safety concerns surrounding drugs like chloramphenicol and telithromycin, which is linked to liver failure^18^, limits our pool of available antibiotics in the face of a rise of multidrug-resistant bacteria. Studying the off-target effect of these drugs can guide the development of safer therapies.

Multiple classes of antibiotics inhibit bacterial translation by binding functionally important sites on the ribosome and disrupting a specific process in translation, resulting in arrested translation on mRNAs at sites characteristic of the mechanism of inhibition^19,20^. In some instances, this relationship is straightforward. For example, the pleuromutilin antibiotic retapamulin binds to the peptidyl-transferase center (PTC) of the bacterial ribosome and impedes translation initiation by interfering with positioning of initiator tRNA^21^, resulting in bacterial ribosomes accumulating at start codons^22^. Inhibition by other antibiotics is more complicated, only disrupting translation when specific sequences are being translated, called context-dependent translation arrest. This is exemplified by chloramphenicol, which simultaneously interacts with the nascent peptide and the peptidyl-transferase center at its binding site and disrupts accommodation of aminoacyl-tRNA, thereby acting as an elongation inhibitor^23^. Due to chloramphenicol’s interaction with the nascent peptide, it selectively causes ribosomes to arrest translation when an alanine, serine, or threonine is in the –1 (penultimate) position of the nascent peptide^24^. Additionally, chloramphenicol does not interfere with the accommodation of glycine due to its minimal size. The mechanisms of these and other antibiotics have been determined by ribosome profiling in bacterial cells. However, it remains unclear whether chloramphenicol or other antibiotics inhibit mitochondrial translation by identical mechanisms.

To determine the sites of translation arrest with, we utilized mitoribosome profiling in HEK293 cells treated with a diverse panel of antibiotics^18,25,26^. Ribosome profiling allows for the mapping of translating ribosomes with codon-level accuracy by identifying mRNA sequences occupied by translating ribosomes. To do this, RNase treatment degrades mRNA unprotected by the ribosome to generate ribosome-protected fragments (RPFs). Because cytoplasmic ribosomes significantly outnumber mitoribosomes, mitoribosomes containing the mitochondrial RPFs (MRPFs) are isolated through sucrose gradient fractionation, where the cytoplasmic ribosome and mitoribosome sediment in the 80S and 55S fractions, respectively (Figure 1A). RPFs are isolated and quantified by next-generation sequencing. Increases in MRPFs reveal the specific codons and sequence contexts prevalent at sites at which mitochondrial translation arrests due to antibiotic treatment.

**Figure 1.**
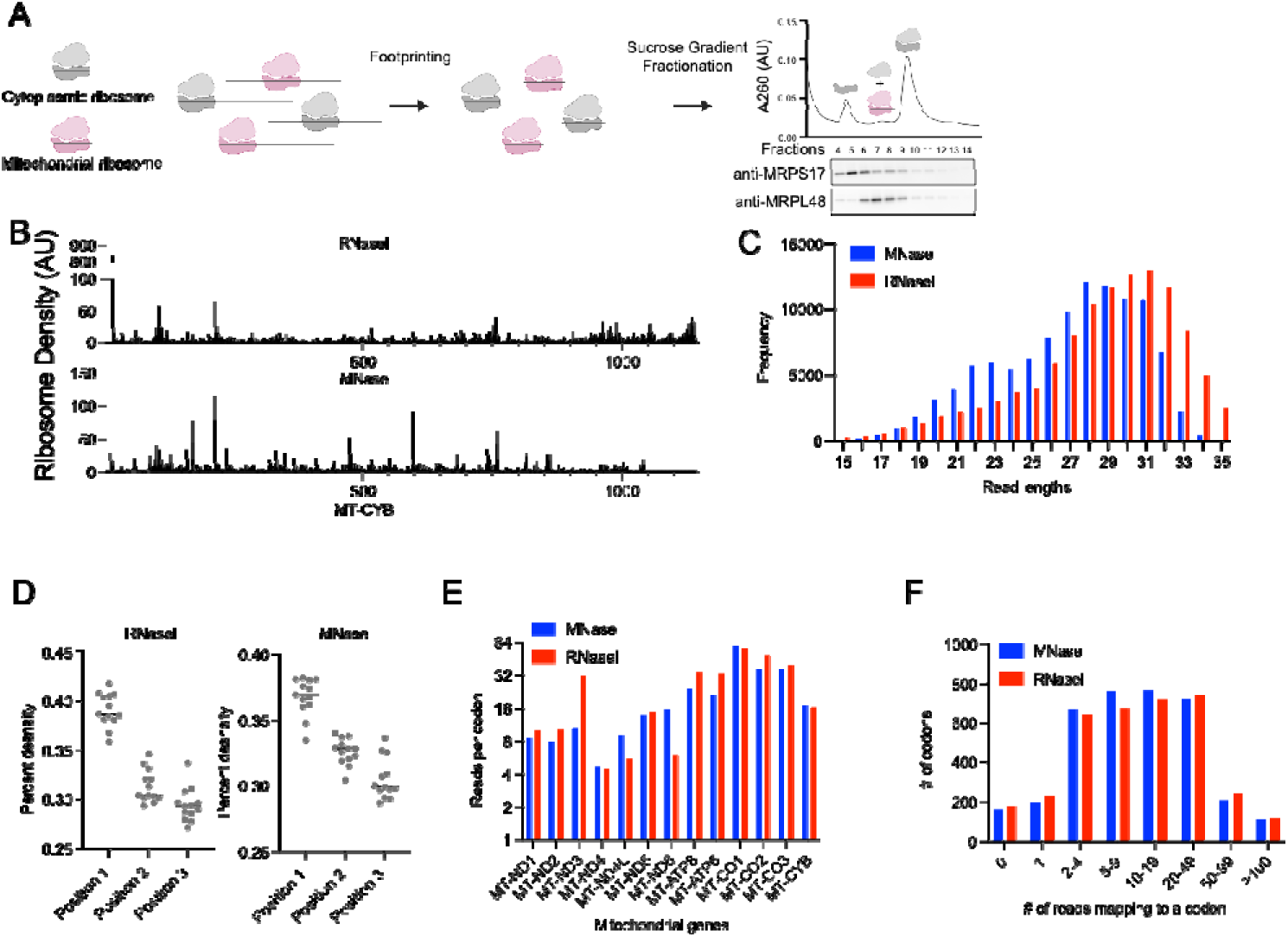
Mitoribosome profiling captures elongating ribosomes. (A) Schematic representation of mitoribosome footprinting and separation from cytoplasmic ribosomes by differential centrifugation. (B) Coverage of MRPFs on *MT-CYB* from mitoribosome profiling generated by either RNase I or MNase footprinting. (C) Distribution of read lengths of MRPFs mapping to the mitochondrial genome. (D) Read phasing analysis of MRPFs, where the 5’ ends of reads are grouped by their subcodon position. Each dot represents an individual mitochondrial gene. (E) Average coverage of mitochondrial genes, calculated by dividing the total number of reads mapping to an ORF and dividing by the length of the ORF. (F) Distribution of mitochondrial codons grouped by the number of mitoribosome P-sites assigned to them.

Understanding the mechanisms of translation inhibitors in mitochondria can guide the development of therapeutics that are more selective to the bacterial ribosome and lays the groundwork for determining how these therapeutics disrupt mitochondrial function. In this study, we utilized mitoribosome profiling in the presence of the bacterial protein synthesis inhibitors retapamulin, tiamulin, josamycin, erythromycin, chloramphenicol, and linezolid to determine their mechanism of action. We found that the mechanism of mitochondrial inhibition is the same as bacterial inhibition for all inhibitors except telithromycin. While telithromycin selectively arrests bacterial ribosomes when either arginine (R) or lysine (K) is in the –1 position of the nascent chain and A-site, called an R/K-X-R/K motif^27^, it instead arrests mitochondrial translation at an R/K/A-X-K motif, suggesting an underlying difference in the mechanism in the two systems. Additionally, nucleotide-level precision of mitoribosome profiling also allowed us to precisely annotate the mitochondrial gene structure. We found that *MT-ND1* and *MT-ND5* genes use alternative translation initiation sites and identified a putative open reading frame on the *MT-RNR1* gene. This work demonstrates that mitoribosome profiling is an effective tool to study the fundamentals of mitochondrial translation and the mechanisms of translation impairment affecting human health.

## Results

### RNase I generates highly phased mitoribosome footprints

To optimize mitoribosome profiling method, we tested two different RNases typically used for ribosome footprinting, RNase I and MNase, followed by the isolation of the 55S mitoribosome by sucrose gradient fractionation. We mapped the sequence reads to the mitochondrial genome and its thirteen open reading frames (ORFs). For each mapped sequence read, we inferred the position of the ribosomal P-site using a 13-nucleotide (nt) offset from the 5’ end, determined by the analysis of footprint distribution surrounding the initiation sites of *MT-ND4*, *MT-ND6*, and the newly reported *MT-ND5-dORF*. We observed coverage across mitochondrial ORFs, suggesting we captured translating mitoribosomes (Figure 1B). Furthermore, analysis of MRPF lengths revealed a distinct difference between RNase I and MNase digested footprint. While both libraries are enriched for reads ∼30 nt in length, MNase footprints showed a bimodal distribution, with a clear secondary peak at ∼22 nts (Figure 1C), which may indicate that MNase-treatment may have captured two distinct conformations of the ribosome. To determine the quality of the ribosome profiling data, we calculated read phasing, which occurs when a nuclease creates uniform MRPFs and the resulting ends of the MRPFs reflect the 3-nt periodicity of elongating ribosomes. While both datasets do show clear phasing, the read phasing is more biased toward position 1 of the codon in the RNase I-treated sample, indicating slightly more consistent digestion (Figure 1D, Table S1). Nevertheless, we observed similar levels of footprint abundance per gene, with the only notable exceptions being that RNase I-digested footprints were enriched on *MT-ND3* and depleted on *MT-ND4L* and *MT-ND6* (Figure 1E). While the cause of the difference is unclear, we did not expect it to interfere when comparing an antibiotic treated sample to a control. Read coverage also appeared similar across the two methods, with 85%-87% codons having between 2-50 reads mapping to them (Figure 1F). Overall, these results indicated that both MNase and RNase I digestion capture mitoribosome footprints with only minor differences. Considering the improved read phasing, we selected RNase I for our subsequent mitoribosome profiling experiments.

### Retapamulin, tiamulin, and josamycin inhibit translation initiation

First, we examined the impact of three antibiotics – retapamulin, josamycin, and tiamulin on inhibiting translation initiation on mitochondrial translation. Retapamulin and tiamulin are members of the pleuromutilin class of antibiotics. They work by binding to the peptidyl transferase center (PTC) of the bacterial ribosome and disrupting initiator tRNA and A site tRNA positioning (Figure 2A, left)^21,22,28–30^. Josamycin is a macrolide antibiotic that binds in the nascent peptide exit tunnel near the PTC and arrests bacterial ribosomes at initiation by interfering with A-site tRNA positioning (Figure 2A, right)^31^. We sought to understand whether these antibiotics also target translation initiation in mitochondria. To this end, we treated HEK293 cells with high concentrations of these antibiotics (100 µg/ml of tiamulin and josamycin, 10 µg/mL of retapamulin) for 30 minutes, performed mitoribosome profiling, and compared the treated experiments to a DMSO treated control.

**Figure 2.**
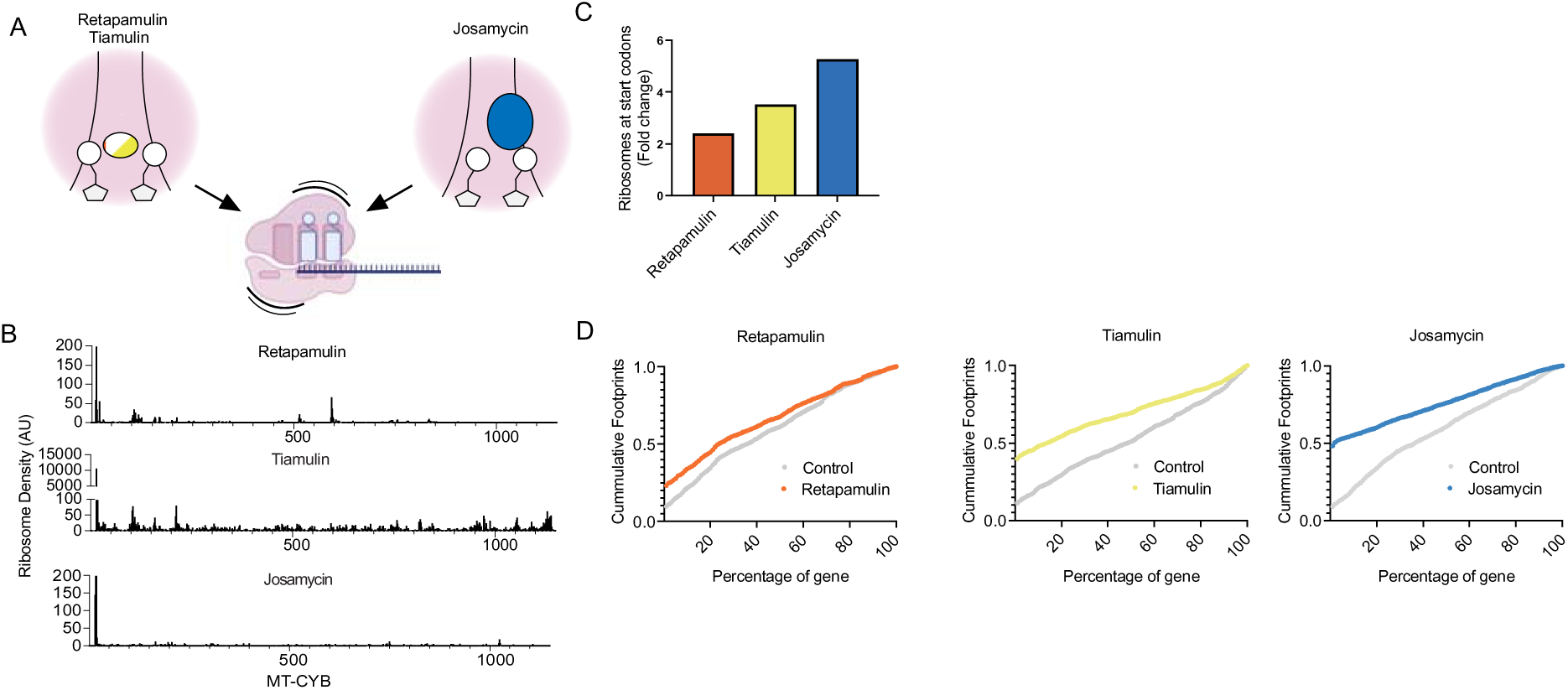
Inhibition of translation initiation in mitochondria. (A) (Left) Schematic representation of retapamulin (orange) or tiamulin (yellow) binding to the PTC and disrupting the first round of elongation. (Right) Josamycin (blue) disrupts accommodation of the first aminoacyl-tRNA. All three inhibit initiation in bacteria. (B) MRPF coverage of *MT-CYB* treated with either retapamulin, tiamulin (note y-axis is segmented), or josamycin. (C) Average fold change increase in the percentage of reads that map to the 5’ of mitochondrial transcripts. (D) Cumulative distribution of ribosome density normalized across all mitochondrial ORFs.

We mapped MRPFs to the 13 mitochondrial genes. On *MT-CYB* and all other genes, we observed that after treatment with antibiotics reads accumulated at the 5’ end of the transcript and MRPFs across the rest of the coding region were lost or decreased (Figure 2B, S1). We quantified the percentage of MRPFs mapping at the 5’ end relative to the total ORF, excluding polycistronic mRNAs. Treatment with translation initiation inhibitors increased the percentage of those reads from 2.4 to 5.2-fold relative to the DMSO-treated cells (Figure 2C). Additionally, we quantified the cumulative footprint abundance moving from the start to the stop codon (Figure 2D). While ribosome footprints are generally evenly spread across the open reading frame in the DMSO control, MRPFs from cells treated with the initiation inhibitors showed a sharp increase in the percentage of reads found at the 5’ end. Together, these results indicate that retapamulin, tiamulin, and josamycin inhibit mitochondrial translation initiation, likely by the same mechanism as in prokaryotes.

### Chloramphenicol, linezolid, and telithromycin arrest mitochondrial translation in a context-dependent manner

Chloramphenicol, linezolid, and telithromycin selectively inhibit bacterial translation when ribosomes are translating specific sequences, in a process called context-dependent translation arrest^24,27^. Linezolid, like chloramphenicol, induces context-dependent arrest when alanine, serine, or threonine are at penultimate position of the nascent peptide (Figure 3A, left). Unlike chloramphenicol and linezolid, telithromycin binds in the nascent peptide exit tunnel near the PTC and adjacent to the nascent peptide^32^. During elongation, telithromycin causes ribosomes to arrest translation at a R/K-X-R/K motif, when the identity of both the penultimate position of the nascent peptide and the incoming aminoacyl-moiety is either arginine or lysine (Figure 3A right).

**Figure 3.**
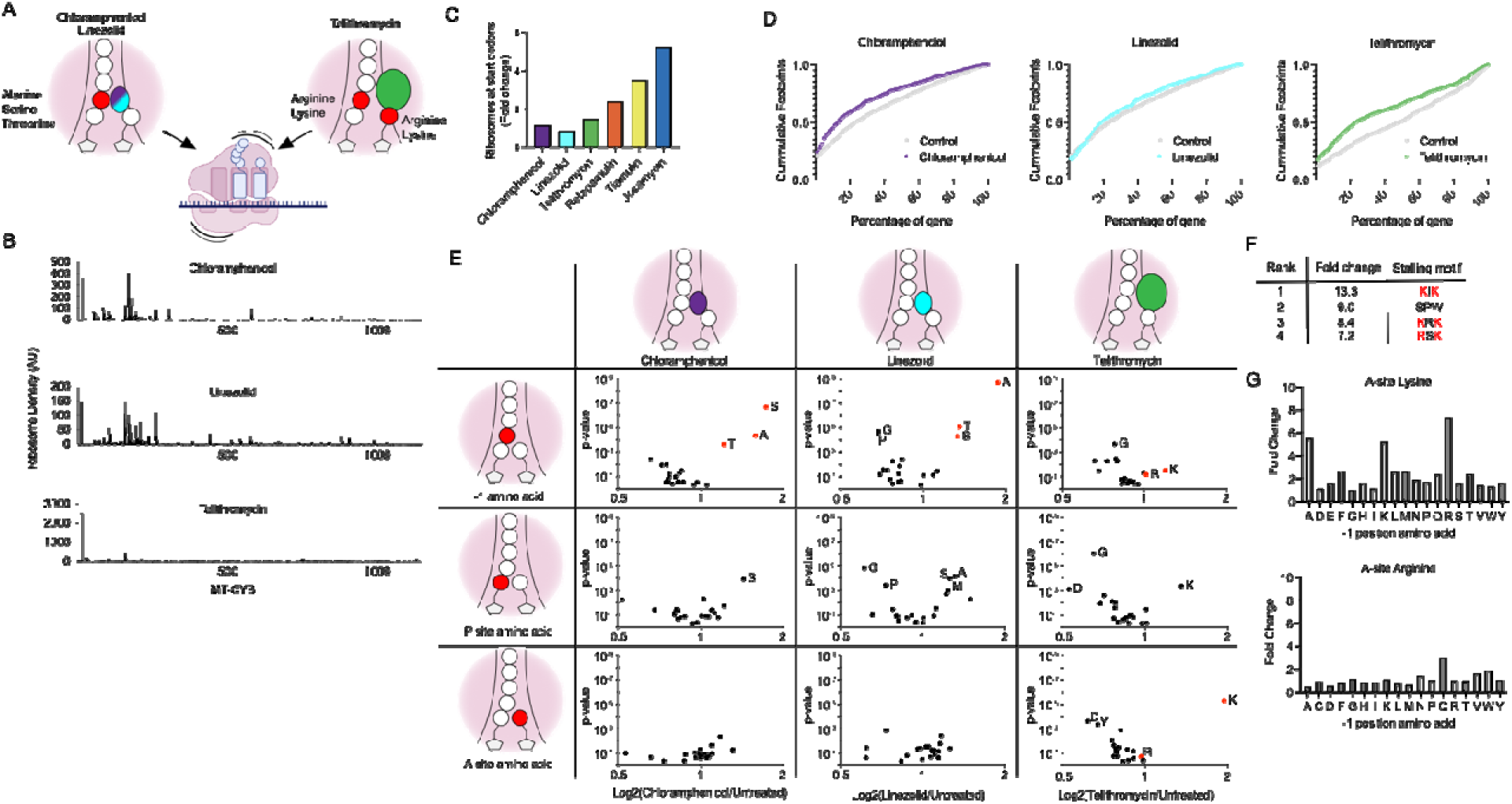
Context-dependent translation arrest by elongation inhibitors. (A) Schematic representation of chloramphenicol (purple), linezolid (cyan), and telithromycin (green) inducing context-dependent translation arrest. (B) MRPF coverage of *MT-CYB* from cells treated with either chloramphenicol, linezolid, or telithromycin. (C) Average fold change increase in the percentage of reads that map to the 5’ of mitochondrial transcripts. (D) Cumulative distribution of ribosome density normalized across all mitochondrial ORFs. (E) Volcano plots of changes in MRPF fold change from cells treated with translation inhibitors. Each dot is the mean log_2_-fold change grouped by the identity of the –1 amino acid (top), P-site amino acid (middle), or A-site amino acid (right). (F) Top 4 strongest stalling sites induced by telithromycin treatment. Red amino acids indicate a classical R/K-X-R/K stalling motif. (G) Fold change in ribosome density, calculated as 2^mean(log2(Telithromycin/Control))^. Bars differ in the identity of the P-site amino acid and contain either a lysine in the A-site (top) or arginine (bottom).

We carried out mitoribosome profiling on HEK293 cells treated with either antibiotic to understand whether these context-dependent translation inhibitors arrest mitoribosomes in the same manner as they arrest bacterial ribosomes. Unlike the cells treated with initiation inhibitors, cells treated with chloramphenicol or linezolid carried ribosome density throughout the entirety of the ORF, suggesting that, as expected, they were not acting as inhibitors of initiation (Figure 3B, S1). Cells treated with telithromycin did show a slight increase in MRPFs abundance at the 5’ end on *MT-CYB.* However, across all genes the levels of MRPFs at the 5’ end was not near the levels seen when using the initiation inhibitor retapamulin and were more similar to chloramphenicol, linezolid, and the DMSO control (Figure 3C). Cumulative distribution of MRPFs suggested that ribosome density was biased towards the 5’ end of the gene for chloramphenicol and telithromycin, but not significantly for linezolid (Figure 3D). These results suggest that these three antibiotics are not acting as initiation inhibitors and that linezolid and telithromycin redistribute MRPFs to the 5’ end of the ORF.

Next, we determined whether this observed redistribution of elongating ribosomes was dependent on the context of the sequence being translated. We calculated the average change of MRPFs depending on the identity of the amino acid at either the -1, P-site, or A-site positions. For both chloramphenicol and linezolid, we observed a specific increase in ribosome footprints when mitoribosomes encountered codons for alanine, serine, or threonine in the -1 position, with a 2-, 2.2-, and 1.3-fold change for chloramphenicol and a 2.3-, 1.4-, and 1.6-fold change for linezolid, respectively (Figure 3E). Additionally, we also observed context-dependent translation arrest induced by telithromycin. Unlike the R/K-X-R/K motif observed in bacteria, we found that telithromycin caused mitoribosomes to arrest at sites with a lysine positioned in the A-site, with an average 2.5-fold increase in MRPFs around those sites, and that neither residue was enriched in the -1 position. Due to the limited size of the mitochondrial genome, there are only nine instances of the R/K-X-R/K motif, nevertheless, when we observed the positions with the greatest fold change, 3 of the top 4 contained the motif with lysine in the A-site (Figure 3F), suggesting that the motif was at least partially recapitulated. To specifically look for the motif we calculated the average fold change when we fixed arginine or lysine in the A-site and varied the identity of the -1 amino acid (Figure 3G). We found that when lysine was in the A-site while either arginine, lysine, or alanine was -1 position, there was strong arrest (> 5-fold change). However, when arginine was in the A-site we did not observe a strong stalling event with any –1 position residue. Therefore, in mitochondria, telithromycin induces stalling at R/K/A-X-K motifs. This motif appears in only 6 of the 13 mitochondrially encoded genes, indicating that synthesis mitochondrial proteins is unequally inhibited (Figure S2 A,B), We concluded that the mechanism underlying context-dependent translation arrest for telithromycin is different in mitochondria compared to bacteria.

### Translation initiation on *MT-ND1 and MT-ND5* occurs at an alternative start codon

During the process of assigning the P-site positions of MRPFs, we observed that no ribosomes were assigned in the first 13 nucleotides of the 5’ end of the transcript. This is not surprising, considering that mitochondrial transcripts often have no or very short 5’ untranslated regions - typically less than 4 nts long - and we applied a 13-nt offset from the 5’ end of MRPFs. It reasons that as mitoribosomes begin translation on a leaderless mRNA, the 5’ end of the resulting MRPF would have a full-length region 3’ of the P-site and no sequence 5’ of the P-site. As the ribosomes move down the transcript, the 5’ end of the MRPF would increase until it is accessible for digestion, at which point it would generate 31 nt-long MRPFs (Figure 4A). As accurate mapping of mitoribosomes at initiation could be a useful tool to understand mitochondrial translation, we attempted to map the location of initiating mitoribosomes at 5’ end using footprint length, rather than the 5’ offset. Because mitoribosome profiling with MNase generated a population of ribosomes with short MRPFs, we hypothesized that these footprints could be indicative of capturing initiating ribosomes (Figure 1C). We plotted the distribution of MRPFs from 20 to 34 nt length that mapped to the 5’ end of the 11 leaderless or short leader ORFs and compared them to reads mapping to the whole ORF. We observed that MRPFs 22-23 nts in length with 6-fold more abundant and MRPFs >28 nts in length were depleted compared to the whole ORF (Figure 4B). As mitochondrial ORFs vary in the length of their 5’ UTR and we expect most ribosomes to be positioned at the start codon, we tested whether leaderless ORFs were enriched in shorter reads compared to ORFs preceded by a short leader. Leaderless ORFs, such as *MT-CO2*, *MT-CO3*, *MT-ND4L* contained MRPFs mainly 20-23 nucleotides in length at the 5’ end (Figure 4C, top). *MT-ATP8*, which contains a start codon beginning at the third nt, contained footprints that were 22-24 nt long and *MT-CO1,* with a start codon at the fourth nt, contained MRPFs 23-24 nt long (Figure 4C, middle). Not only did we observe longer MRPFs in short leader ORFs, the size of the MRPFs also appeared proportional to the location of the start codon. Together, these results indicate that the size of the footprint is directly related to the start codon position at the 5’ end of mitochondrial mRNAs.

**Figure 4.**
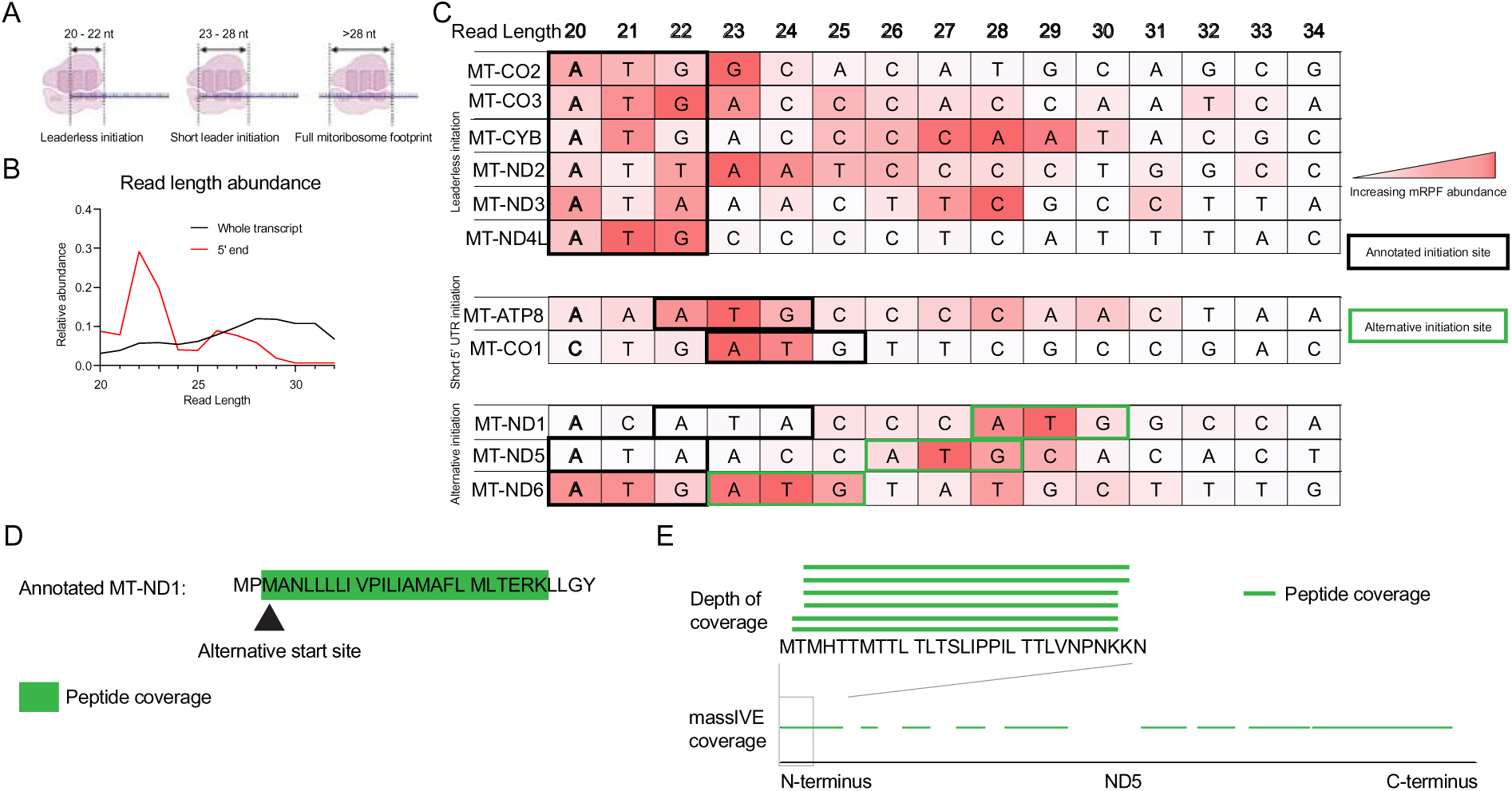
MRPF sizes indicate use of alternative initiation sites. (A) Schematic representation of changing MRPF sizes as a function of the mitoribosome’s position near the 3’ end of mRNAs. (B) Read length measurement of reads mapping to the 5’ ends of mitochondrial transcripts (red) compared to the entire ORF (black). (C) Read lengths of MRPFs mapping to the 5’ ends of mitochondrial transcripts. Each row contains the nucleotide sequence of each transcript starting from the 5’ end. The coloring of the box is proportional abundance of MRPFs with read lengths from 20-34 nts. Nucleotide sequence and read abundance are overlayed to compare the abundance of reads to potential ribosome start sites. Transcripts are grouped by being leaderless, having a leader, or having a potential alternative start site. Black boxes indicate annotated start codons and green boxes indicate alternative initiation sites. (D) Visual representation of the peptide coverage (green bars or highlights) from mass spectrometry data covering the N-terminus of *MT-ND1* or (E) *MT-ND5*.

Two genes, *MT-ND*1 and *MT-ND5,* deviated from this trend. The annotated start site for *MT-ND5* is a leaderless AUA codon (a start codon in mitochondria) but we observed few 20-22 nt MRPFs (Figure 4C, bottom). Similarly, the annotated start site for *MT-ND1* is also an AUA codon that begins at the third nt and it is depleted in reads for MRPFs that are smaller than 25 nts. Instead, MRPFs derived from the 5’ end of both genes are enriched in longer reads: 27 nts-long for *MT-ND5* and 28-29 nts-long for *MT-ND1*. Curiously, the third codon for both genes is an in-frame AUG and the read lengths proportionally correspond to mitoribosomes positioned at these start codons, suggesting mitoribosomes initiating translation at these sites. MRPFs generated from cells treated with initiation inhibitors and footprinted by MNase yield similarly sized reads at the 5’ end of *MT-ND1* and *MT-ND5* (Figure S3). Using protein mass spectrometry, we tested whether peptides matching the N-terminus of these proteins would correspond to initiation at the canonical start codons or alternative start codons. We identified a peptide that mapped specifically to the truncated version of MT-ND1 and not the full-length peptide, indicating that translation can initiate from the alternative start codon (Figure 4D). While we did not identify N-terminal peptides mapping to MT-ND5 from our mass spectrometry data, the protein mass spectrometry database MassIVE^31^ covered the N-terminal region of MT-ND5 (Figure 4E). Consistent with our mitoribosome profiling data, the majority of N-terminal fragments did not include the amino acids from the annotated start or second codon but began with methionine at codon 3. We also observed that MRPFs on *MT-ND6,* which initiates at the first of two AUGs codon immediately at the 5’ end, correlated to ribosomes positioned at both AUG codons, indicating that this gene may initiate translation from the second AUG codon as well. However, we did not observe any peptides mapping to the N-terminus of *MT-ND6*. Together, this data shows that *MT-ND1* and *MT-ND5* may not initiate from their canonical AUA codons but may use alternative in-frame AUG codons. Additionally, our findings indicate that the mapping of mitochondrial translation initiation events by size, rather than 5’ offset is an accurate strategy.

### Mitoribosome profiling may reveal novel translation events

During our analysis of 5’ end mapping, we did not remove reads mapping to non-coding RNAs (ncRNAs) before mapping to the mitochondrial genome and performing P-site assignment. Consequently, we observed that some tRNAs and rRNAs had an abundance of reads mapping to 13 nts away from their 5’ ends, similarly to what we observe on mRNAs for translation initiation. In addition, some of these ncRNAs contained a potential start codon near the 5’ end. These reads could either be products of random digestion of the abundant background of ncRNAs or be genuine MRPFs. We hypothesized that if these reads were the result of mitoribosomes initiating or attempting to initiate translation at the 5’ end of the ncRNAs, then we could use the same analysis of read length distribution at 5’ ends of mRNAs we performed above and determine of the sizes correlated to an AUG, AUU, GUG, or UUG as potential start codons. While GUG and UUG are not used in canonical mitochondrial initiation they were included in the search because noncanonical initiation has been observed at GUG^33^ and both are frequently utilized in bacterial initiation. We identified sites on 4 tRNAs that contained MRPFs with a size that correlated to either AUG or UUG codons (Figure 5A). The predicted ORFs and thus resulting potential translation events varied widely. On *MT-TF*, the start codon is followed by an immediate stop codon. The ORF on *MT-TI* is 3 codons long and terminates with the non-canonical mitochondrial termination codon AGA. On *MT-TY, a* 7-codon ORF is terminated by a UAA codon. *MT-TS1* contains the largest open reading frame identified on tRNAs, at 20 codons and it terminates with at the U AGA sequence, as used in *MT-CO1*^34^. Additionally, we identified a putative start site at the 5’ end of *MT-RNR1* (Figure 5A) and mitoribosome profiling of cells treated with initiation inhibitors and digested with MNase show similar footprint sizes (Figure S3). However, this site did not occur at a canonical AUG or AUU codon, but instead at a UUG codon. This putative start codon is followed by a 58-codon ORF and terminates at a UAG stop codon.

**Figure 5.**
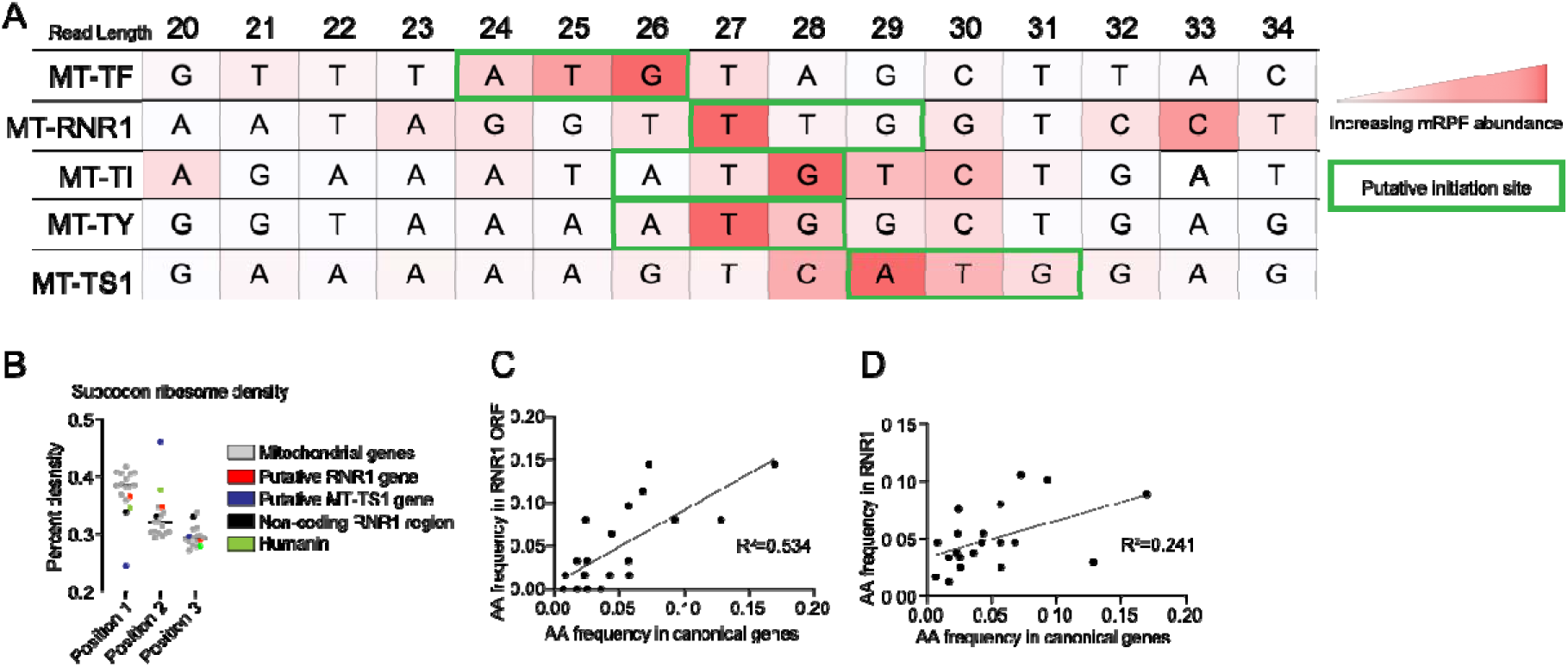
MRPFs on ncRNAs indicate potential novel ORFs. (A) Read lengths of MRPFs mapping to the 5’ end of ncRNAs. Each row contains the nucleotide sequence beginning from the 5’ end of listed ncRNAs. The coloring of the box is proportional the abundance of MRPFs with read lengths 20-34 nts. The nucleotide sequence and read MRPF abundance are overlayed to identify potential new start sites. Putative start sites are indicated by green boxes. (B) Read phasing analysis of all canonical ORFs (grey), a mutative ORF located on *MT-RNR1,* the non-coding region of *MT-RNR1,* and Humanin. (C) A correlation plot comparing the frequence of amino acid usage in canonical ORFs compared to the putative ORF on *MT-RNR1* or (D) the non-coding region of *MT-RNR1*.

To determine whether translation elongation occurs at these putative ORFs, we determined the read phasing characteristic for MRPFs derived from translating ribosomes at these ORFs. However, the sites on *MT-TF*, *MT-TI*, and *MT-TY* are short and close to the 5’ end, hindering phasing analysis and thus do not allow us to conclude whether our sequence reads are indicative of translation. While the ORF on *MT-TS1* is longer, MRPF density was low and we did not observe read phasing and thus is likely not translated (not shown). The reads mapping to the ORF on *MT-RNR1* had a phasing profile similar to those mapping to the 13 canonical mitochondrial genes (Figure 5B). In contrast, reads mapping to the remainder of *MT-RNR1* or mapping to the region of *MT-*RNR2 that codes for the putative mitochondrial micropeptide Humanin, shows no or conflicting read phasing. When compared to the amino acid composition of the mitochondrially-encoded proteome, the amino acid composition of the resulting peptide shows a stronger positive correlation (R^2^=0.534) (Figure 5C) to the remaining non-coding portion of *MT-RNR1* (R^2^=0.241) (Figure 5D). Although mitoribosome profiling suggests that this putative ORF is translated, we did not detect any corresponding peptides in the protein mass spectrometry of isolated mitochondria. Nevertheless, our mass spectrometry dataset, or the MASSIVE database do not cover many regions of the mitochondrial proteome and thus, the absence of a spectrum is not necessarily evidence that the peptide is not translated. The read phasing analysis suggests this region of *MT-RNR1* is translated and could encode a 15^th^ mitochondrial protein. Overall, this data shows that our mitoribosome profiling can be used as a tool to study non-canonical translation events resulting from ncRNAs.

## Discussion

In this work, we sought to understand the mechanisms of bacterial protein synthesis inhibitors on their primary off-target, the mitoribosome. We found that inhibitors of bacterial translation initiation, retapamulin, tiamulin, and josamycin, also act as initiation inhibitors in mitochondria. Likewise, chloramphenicol and linezolid selectively inhibit elongation when the mitoribosome carries either an alanine, serine, or threonine in the penultimate position of the nascent peptide, matching their mechanism in bacteria. In contrast, the context-dependence of telithromycin differed when arresting mitoribosomes, wherein it preferentially arrested translation at R/K/A-X-K motif.

Context-dependent translation arrest by chloramphenicol and linezolid suggests similarities between binding of the bacterial ribosome and the mitoribosome. Structural and biochemical studies have demonstrated that binding of chloramphenicol and linezolid is stabilized by alanine, serine, or threonine in the penultimate position of nascent peptide^35,36^. Our findings suggest a similar or identical interaction is occurring in the mitoribosome. In contrast, these inhibitors are unable to inhibit peptide bond formation in bacteria when glycine is positioned in the A-site, due to the minimal size of the side chain, allowing the inhibitor and the aminoacyl-moiety to coexist in the PTC. We did not observe a lack of mitoribosome translation arrest when glycine was positioned in the A-site. This may have technical reasons, e.g. it may be caused by an insufficient depth in sequencing or indicate that the relatively small mitochondrial genome limits the variety of stalling and non-stalling events we can observe. But more likely it shows that chloramphenicol and linezolid are capable of arresting translation when glycine is in the A-site. Recently, mitoribosome profiling analysis showed that chloramphenicol added during lysis did not induce context-dependent translation arrest^37^. This suggests that during lysis, there is insufficient mitochondrial translation to induce a redistribution of mitoribosomes rather than an inability of the chloramphenicol to induce context-dependent arrest.

During preparation of this manuscript, two other groups studied the effect of translation inhibitors on mitochondrial translation. One group’s mitoribosome profiling in the presence of chloramphenicol and linezolid agreed with our observed context-dependent translation arrest^38^. Their structure of linezolid in the mitochondrial peptidyl-transferase center also suggested some nuanced differences in binding between the bacterial and the mitochondrial ribosome. Another group carried out mitoribosome profiling in the presence of retapamulin^37^. However, they observed additional ribosome density throughout the coding region and potentially some associated with putative start codons. This differs significantly from our observed accumulation at only annotated start sites and may be due to differences in mitoribosome profiling methods. Future work may reveal how each strategy contributes to our understanding of mitochondrial translation.

Inhibition of mitochondrial translation through either small molecules or mutations localized in mitochondrial translation factors can induce a wide range of pathologies^39–41^. However, it is unclear how or whether the relationships between the mechanism of translation inhibition relates to the resulting disease. Recently the proteins mtRF-R and MTRES1 were shown to act as quality control factors, rescuing stalled mitoribosomes by removing the nascent peptide and peptidyl-tRNA^42^. As the mitoribosome can arrest translation through a variety of mechanisms and small molecules, it remains unclear whether these factors can rescue all stalled mitoribosomes. Further work to dissect the relationship between mechanisms of inhibition, activation of rescue factors, and stress response can complement our understanding of pathologies resulting from mitoribosomopathies and antibiotic treatment.

In this study we also reported that mitoribosomes accumulate at in-frame AUG codons shortly downstream of their annotated initiation sites on two mitochondrial genes, *MT-ND1* and *MT-ND5*. The biological function of these events and the start codon preference between canonical and alternative sites is unclear. Other work does not preclude the usage of these codons. While mass spectrometry studies of mitochondrial proteins derived from bovine do show full length peptides^43^, due to differences in AUA and AUG codon usages at these sites, the accumulation of mitoribosomes at the 3^rd^ codon may not occur. A recent structural study of Complex I purified from HEK293T cells models the N-terminal residues for all mitochondrially-encoded subunits except for the first two residues of *MT-ND1* and *MT-ND5*, consistent with our observations^44^. This may be due alternative start codon usage or difficulty resolving the resides on the surface of Complex I. Others have noted that haplogroups L1b and F2 contain a single nucleotide polymorphism at either position T3308C or T12338C respectively, which disrupts the annotated AUA start codons of *MT-ND1* and *MT-ND5* yet do not result in any discernable pathogenicity^45–47^. The authors noted that the same AUG codons we identified as start codons could be used in lieu of the canonical AUA start sites. Because we observe MRPF lengths primarily correlating with the AUG codons, our results agree with a model wherein a downstream start codon may serve as the alternative start sites for these genes. Alternative to their function as start codons, the accumulation of MRPFs at these codons may have some unknown role in translation regulation or programmed pausing.

Recently, mitoribosome profiling was used to identify an open reading frame in region immediately 3’ of the gene *MT-ND5,* which we confirmed observing in our profiling experiments^48^. However, this uses an internal initiation mechanism for polycistronic genes, similar to *MT-ND4* and *MT-ATP6.* Until now, accurate mapping of mitochondrial translation events at the 5’ ends of transcripts has been unavailable. We identified MRPFs that map to putative start codons occurring on mitochondrial MT-tRNAs and *MT-RNR1*, based on the size of the reads occurring at the 5’ end of the RNA and their relationship to a start codon localized in the sequence. The site of *MT-RNR1* was further supported by analysis of the resulting ORF, suggesting translation initiation and elongation occurs. Curiously, tRNA^Tyr^, encoded by *MT-TY*, is polyadenylated during processing, which may be related to our identification of a ribosome binding event on this tRNA^49^. Due to both the short ORFs, minimal read coverage, and lack of a detectable peptide we could not determine if translation elongation occurred on the mitochondrial tRNAs. These sites may be unproductive mitoribosome binding events or simply from tRNAs partially digesting during MNase treatment. Future work is needed to understand whether these are actual translation events and to determine the function of these potential novel peptides. Together our findings improve our understanding of mitochondrial translation initiation, gene expression, and the mechanisms of translation inhibition.

## Materials and Methods

### Isolation of mitoribosomes for profiling and western blotting

Mitoribosome profiling was carried out in HEK293 Flp-In T-REx cells. 15 cm plates were treated with either DMSO as a control or 100 µg/mL of protein synthesis inhibitor for 1 hour, washed with 4°C PBS, and flash frozen by floating the plate on liquid nitrogen. Cells were collected by scrapping on ice with 600 µL of lysis buffer (1.5x Lysis buffer (30 mM Tris–HCl pH 7.8, 150 mM KCl, 15 mM MgCl_2_, 1.5 mM DTT, 1.5% Triton X-100, 0.15% NP40, 1 × complete phosphatase and protease inhibitors)) and lysed by passing through a 27-G needle 10 times. Lysate was centrifuged for 10 minutes at 21,000 g at 4°C and the supernatant was collected.

300 µL of lysate was footprinted with either digested with 1,5000 U of MNase, 10 µL of SUPERase-In, and 5 mM of CaCl_2_ at 22°C for 1 hour and stopped by adding EGTA to a concentration of 6 mM or with 8 U/µL of RNase I at 22°C for 1 hour and stopped by adding 10 µL of SUPERase-In. Titration of the digestion enzymes was not performed and is based on previous published work^26,48^. The digestion was clarified by centrifugation at 21,000 g for 5 minutes. Lysates were loaded onto a 5%-30% sucrose gradient (20 mM Tris-HCl pH 7.8, 100 mM KCl, 10mM MgCl_2_, 1 mM DTT) and spun for 2.5 hours at 40,000 rpm in a SW40 rotor. 55S mitoribosome fractions were collected using a BioComp Fractionator and MRPFs were isolated by SDS/hot phenol/chloroform extraction.

### Mitoribosome profiling library construction

Samples were run on a 15% TBE-Urea polyacrylamide gel and footprint sizes from 15-34 nts were excised and footprints were eluted in 0.3 M NaOAc, 1 mM EDTA, and 0.25% SDS followed by purification by Oligo Clean & Concentrator. Samples were treated with Quick CIP. 3’ adapters (5’-rAppNN-6 nt UMI-TGGAATTCTCGGGTGCCAAGG-L) were ligated using Rnl2(1-249)K227Q ligase for 2 hours at 16°C and pooled. Pooled samples were treated with T4 PNK for 30 minutes at 37°C. A 5’ chimeric DNA-RNA adapter (GTTCAGAGTTCTACAGTCCGACGATCrNrNrNrN) was ligated to the samples with T4 RNL1. Reverse transcription was carried out using SuperScript IV followed by PCR amplification by Taq DNA polymerase. Libraries were size selected using a 3% agarose Pippin prep and final libraries were PCR amplified. Samples were sequenced as either singe- or paired-end 50 nt reads on either a nextseq550 or novaseq.

### Mitoribosome profiling analysis

Adapter sequences were removed with cutadapt and aligned with bowtie2 to a fasta containing rRNA and tRNA sequences. To identify stalling sites, reads were mapped to a custom mitochondrial transcriptome containing all 5’ UTRs, coding sequences, and 3’ UTRs for all processed mRNAs. Reads were aligned with bowtie2. A 13 nt P-site offset was assigned using Plastid and mapping reads relative to the start codons of *ND4*, *ATP6*, *MT-ND5-dORF.* We determined the relative ribosome density by normalizing the number of reads at each codon by the total number of reads mapping to its respective gene for each sample. Fold change was then calculated by dividing the relative ribosome density in a treated sample to an untreated sample, excluding sites that carried less that 0.1% of the total ribosome density of each gene. To identify putative novel translation initiation sites, we mapped our footprints to the entire mitochondrial genome and visually inspected the 3’ ends of mitochondrial tRNAs and rRNAs.

### Mass spectrometry analysis of mitochondrial proteins

Mitochondria were isolate from 2×10^7^ HEK293 cells using the Mitochondria Isolation Kit for Cultured Cells (Thermo Scientific). Mitochondrial proteins were processed for trypsin/LysC digestion and the digested peptide mixture was then concentrated and desalted using C18 column. Reconstituted desalted peptides in 25 μl of 0.1% formic acid. 12 μl of peptides from each sample was analyzed by 110 min LC/MS/MS run. The LC/MS/MS analysis of tryptic peptides for each sample was performed sequentially with a blank run between each two sample runs using a Thermo Scientific Orbitrap Exploris 240 Mass Spectrometer and a Thermo Dionex UltiMate 3000 RSLCnano System. Peptides from trypsin digestion were loaded onto a peptide trap cartridge at a flow rate of 5 μL/min. The trapped peptides were eluted onto a reversed-phase Easy-Spray Column PepMap RSLC, C18, 2 μM, 100A, 75 μm × 250 mm (Thermo Scientific) using a linear gradient of acetonitrile (3-36%) in 0.1% formic acid. The elution duration was 60 min at a flow rate of 0.3 μl/min. Eluted peptides from the Easy-Spray column were ionized and sprayed into the mass spectrometer, using a Nano Easy-Spray Ion Source (Thermo) under the following settings: spray voltage, 1.6 kV, Capillary temperature, 275°C. Other settings were empirically determined. Raw data files were searched against human protein sequences database and client target protein sequences database using the Proteome Discoverer 2.5 software (Thermo, San Jose, CA) based on the SEQUEST algorithm. Carbamidomethylation (+57.021 Da) of cysteines was set as fixed modification, and Oxidation / +15.995 Da (M), Phospho / +79.966 Da (S, T, Y), and Deamidated / +0.984 Da (N, Q) were set as dynamic modifications. The minimum peptide length was specified to be five amino acids. The precursor mass tolerance was set to 15 ppm, whereas fragment mass tolerance was set to 0.05 Da. The maximum false peptide discovery rate was specified as 0.01. The resulting Proteome Discoverer Report contains all assembled proteins with peptides sequences and peptide spectrum match counts (PSM#) and MS1 peak area relative abundance.

## Data sharing

Mitoribosome fastq files, wig files mapping to mitochondrial mRNAs, and read length files for MNase treated samples mapping to chrM are available through GEO Series GSE277563.

## Author contributions

J.M. Conceived the project, designed and executed experiments, analyzed data, and wrote the manuscript. E.Y. Analyzed mitoribosome profiling data. M.H. supervised the project, advised on data interpretation and experimental design, and edited the manuscript.

## Author contributions

We’d like to thank Nicholas Guydosh (NIDDK), Sezen Meydan (Vanderbilt University), Astrid Haase (NIDDK), the Haase lab members, and the Hafner lab members for their feedback on this work. We thank Daniel Ben Halvey (Tel Aviv University) for his assistance in carrying out ribosome profiling and Sarah Young-Baird (Uniformed Services University) for assistance with sucrose gradient fractionation. We thank Stefania Dell’Orso and Faiza Naz (NIH/NIAMS Genomic Technology Section) for sequencing experiments.

## Supplementary Figures and Tables

**Table S1.**
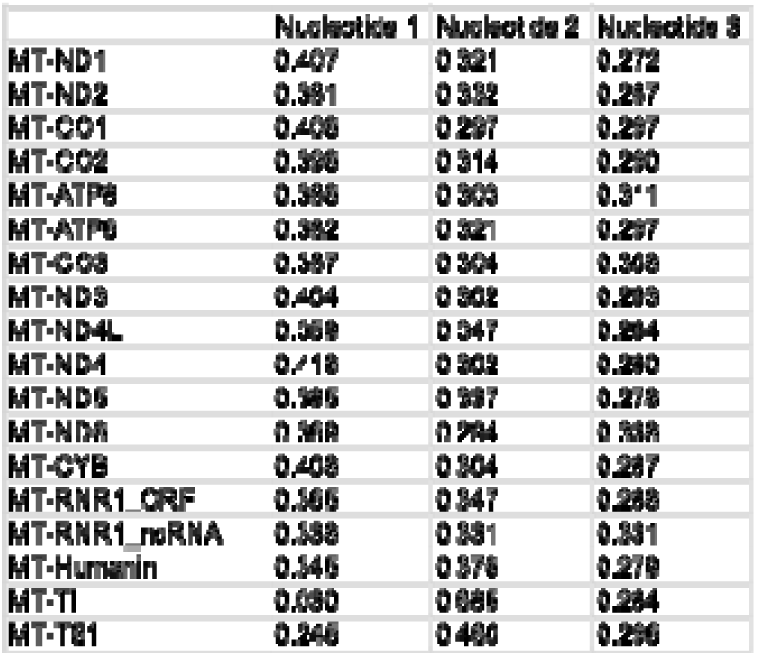
Read phasing values for mitoribosomes footprinted with RNaseI. The read phasing values for the canonical and putative ORFs.

**Figure S1.**
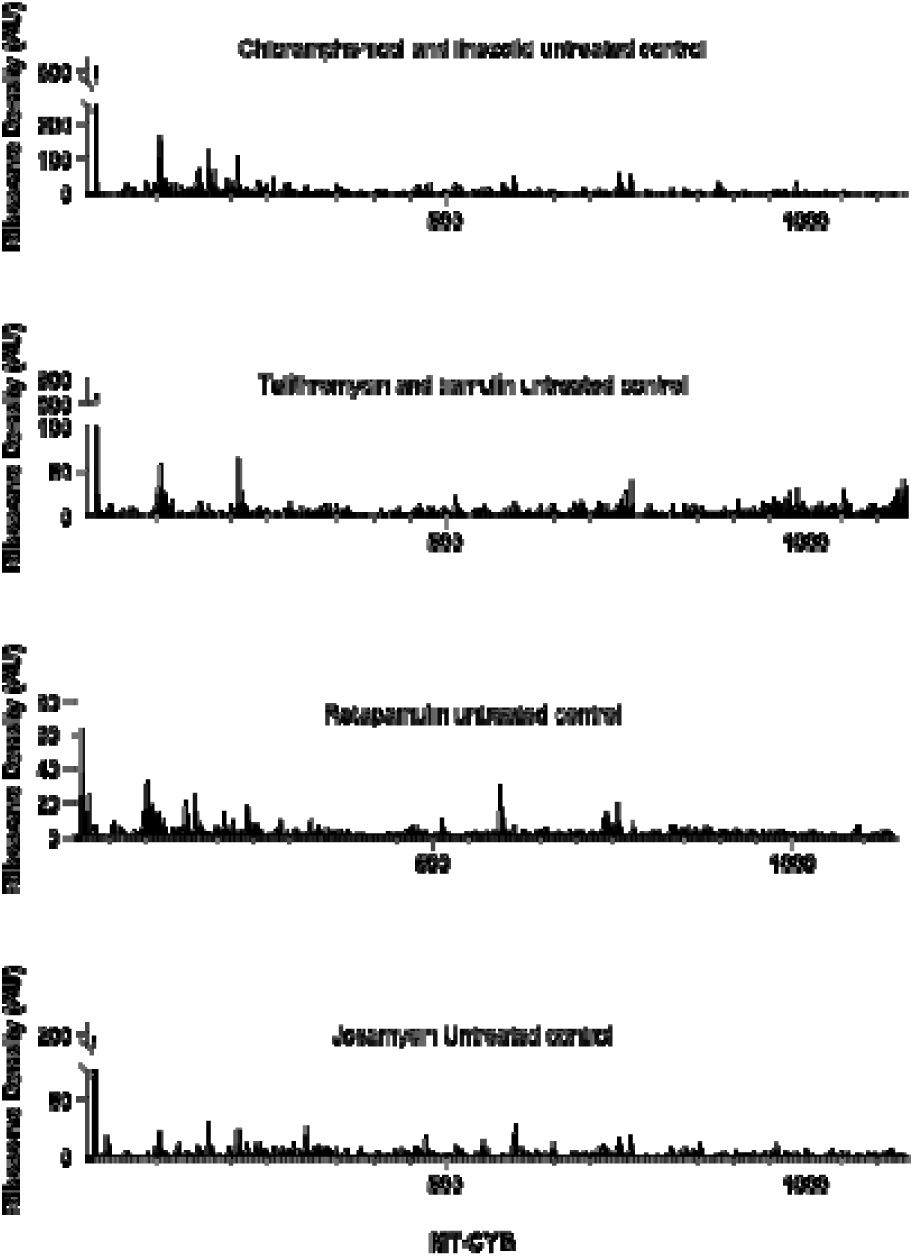
Mitoribosome profiling traces of *MT-CYB*. Individual traces from mitoribosome profiling from each untreated control labeled with its’ corresponding antibiotic-treated sample.

**Figure S2.**
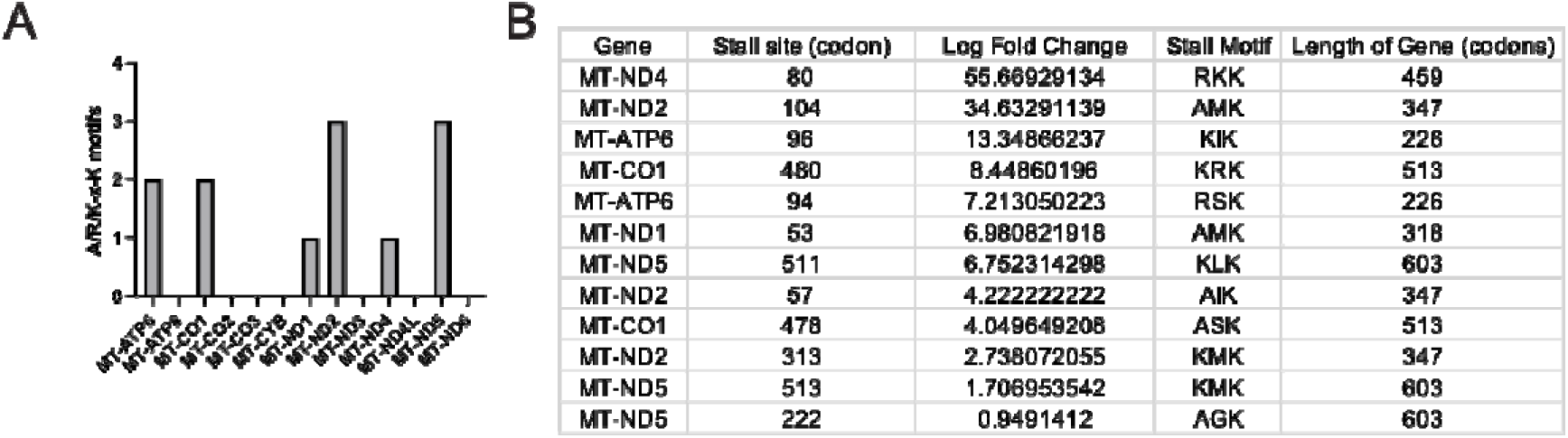
Distribution of A/R/K-X-K telithromycin stalling site. (A) The frequency of each telithromycin stalling sites across canonical mtDNA-encoded genes and (B) the measured log fold change at those sites as compared to an untreated control.

**Figure S3.**
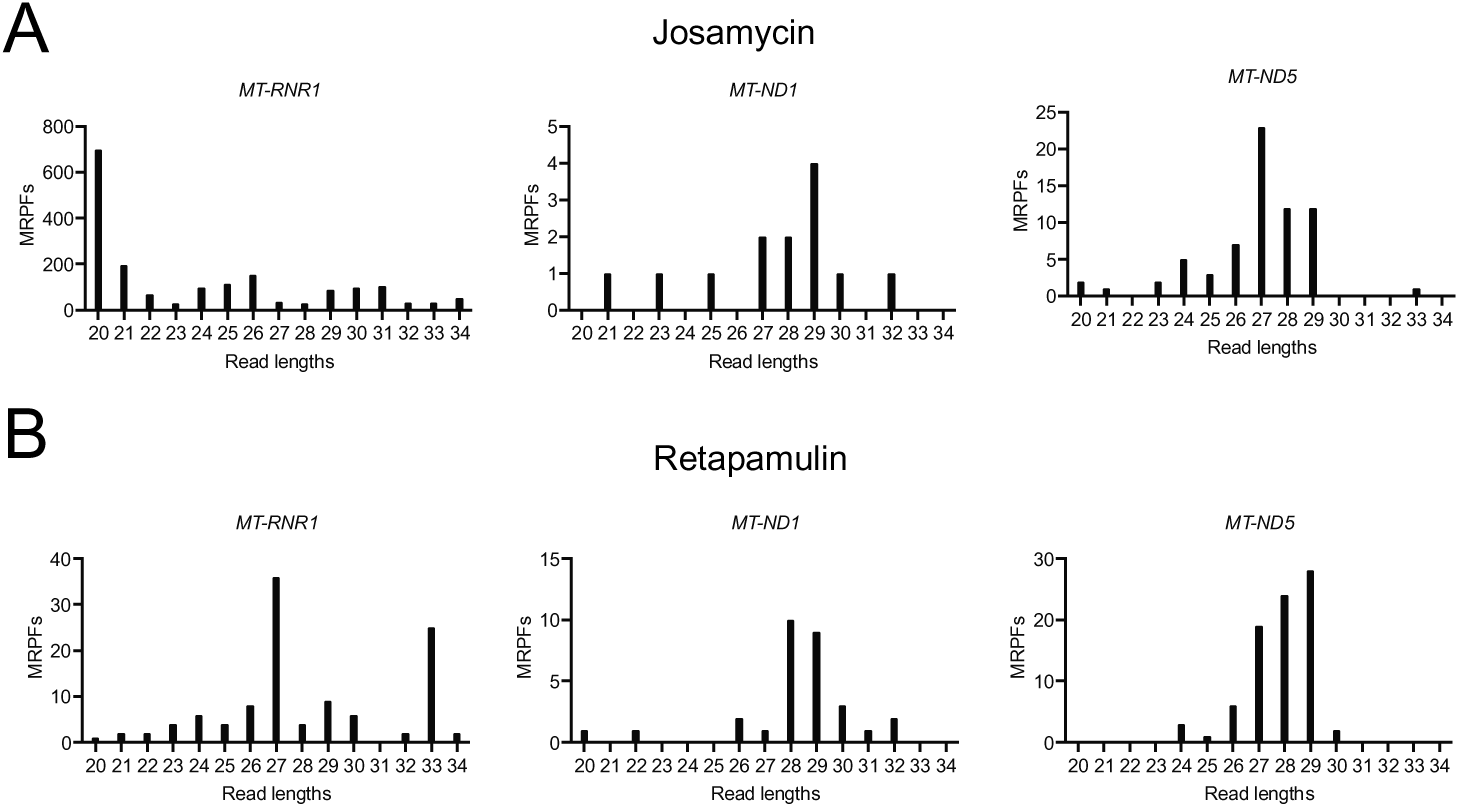
MRPF lengths after treatment with initiation inhibitors correlate to alternative initiation sites. (A) Read lengths of MNase-generated MRPFs from cells treated with josamycin or (B) retapamulin.

